# A broad-host-range *Rhizobium rhizogenes* strain enables transient expression across diverse crops and establishes functional assays in faba bean

**DOI:** 10.64898/2026.04.20.719558

**Authors:** Freddie King, Juan Carlos Lopez-Agudelo, Christopher Stephens, May Htet Aung, Tarhan Ibrahim, Enoch Lok Him Yuen, Ao Chen, Nick Eilmann, Saskia Jenkins, Cristina Vuolo, Yueh-Ning Swee, Wei-Jia Liu, Samuel Bruty, AmirAli Toghani, Chih-Horng Kuo, Erh-Min Lai, Jiorgos Kourelis, Lida Derevnina, Chih-Hang Wu, Tolga Bozkurt

## Abstract

*Agrobacterium*-mediated transient expression has revolutionized plant research, enabling numerous landmark discoveries across diverse areas of plant biology. Yet this powerful approach remains largely confined to solanaceous species, leaving most economically important crop families without a comparable rapid assay platform. Here, we show that an engineered *Rhizobium rhizogenes* strain, AS109, mediates efficient transient expression across diverse dicot species spanning multiple taxonomic families, consistently outperforming commonly used laboratory agrobacterial strains. Leveraging the broad host range of AS109, we establish a suite of functional assays in faba bean (*Vicia faba*), including protein localisation, RNA interference-mediated gene silencing, cell-surface elicitor recognition screens, nucleotide-binding leucine-rich repeat receptor (NLR) activation, and infection cell biology at the host-pathogen interface. We further demonstrate that both singleton NLRs and sensor-helper NLR pairs from Solanaceae retain effector recognition and cell death activity when transferred into faba bean, establishing a rapid platform for evaluating cross-family transferability of disease-resistance genes. AS109 thus provides an accessible and versatile chassis for functional genomics in non-model crops, bridging the widening gap between hypothesis generation and experimental validation across diverse plant species.

## Introduction

Advances in genomics and computational biology, including protein structure prediction and large-scale genomic analyses, are rapidly accelerating hypothesis generation in the plant sciences. Yet the capacity to experimentally test these hypotheses has not kept pace, as biological assays remain slow, labour-intensive, and often restricted to a handful of model species. *Agrobacterium*-mediated transient expression, particularly in *Nicotiana benthamiana*, has been a major solution to this experimental bottleneck, enabling rapid experimental validation and landmark discoveries in plant-pathogen interactions, effector biology, immune receptor discovery and function, structural biology, and gene silencing pathways such as RNA interference (RNAi) and virus-induced gene silencing (Lawson et al., 2026; Yang and Xu, 2026; Bally et al., 2018). These achievements underscore the transformative power of transient assays for functional studies in plants. However, transient expression remains largely confined to solanaceous species (Wroblewski et al., 2005), leaving many economically important crop groups, including legumes, brassicas, and Asteraceae without a comparable experimental platform.

Recent efforts to enhance *Agrobacterium*-mediated transient expression have focused on rewiring virulence gene regulation, optimizing plasmid architecture, suppressing host immunity, and refining infiltration conditions (Matsuda, 2026; Goralogia et al., 2025; Aliu et al., 2024; Beritza et al., 2024). While these strategies improve expression levels, a fundamental limitation persists: most transient expression assays rely on *Agrobacterium fabrum* (formerly *A. tumefaciens*) C58-derived strains, whose narrow host range excludes the majority of crop species. We recently developed *Rhizobium rhizogenes* strain AS109 (referred to as AS109 hereafter) as a new platform for transient expression in solanaceous crops (Lopez-Agudelo et al., 2025). Given its distinct genetic background and delivery properties, we asked whether AS109 might support transient expression beyond solanaceous hosts. Here, we show that AS109 exhibits a broad host range spanning diverse dicot families. Given the global importance of legume crops for dietary protein, and the limited transient systems in this family, we sought to use AS109-mediated expression to establish a new model species in Fabaceae. To this end, we use AS109 to build a transient expression system in faba bean (*Vicia faba*), enabling protein localization, gene silencing, immune receptor assays and infection cell biology. Together, these findings position AS109 as a versatile transient-expression chassis for non-solanaceous crops and provide a practical framework for rapid functional testing of candidate genes and pathways emerging from genomic and computational discovery.

## Results

### *Rhizobium rhizogenes* AS109 enables transient expression across diverse dicot crops

To assess the applicability of AS109 across diverse plant species, we compared AS109 with several commonly used laboratory *Agrobacterium* strains (EHA105, LBA4404, GV3101::pMP90, AGL-1, and C58C1::pTiB6S3ΔT) in 30 plant species representing multiple taxonomic groups and families (Fig. 1a,b, Supplementary Figs. 1-4 and Supplementary Table 1). We introduced bacterial suspensions into leaves by simple syringe infiltration, and assessed transient expression efficiency using the RUBY reporter, which encodes three enzymes that convert tyrosine into red-purple betalain pigments, enabling direct visual observation and quantitative analysis (He et al., 2020). We first focused on species belonging to Solanaceae, in which AS109 was previously shown to be effective (Lopez-Agudelo et al., 2025). Consistent with our previous report, AS109 showed superior transient expression efficiency compared with other tested laboratory strains on petunia (*Petunia hybrida*), tobacco (*N. tabacum*), potato (*Solanum tuberosum*), black nightshade (*Solanum nigrum*), goji berry (*Lycium barbarum*), and goldenberry (*Physalis peruviana*) (Fig. 1c and Supplementary Fig. 1c). Although *N. tabacum* is widely used for transient expression assays, transformation efficiency can vary substantially between leaves. Notably, while all other tested laboratory strains exhibit poor transient expression in older (lower) leaves, AS109 mediated highly reliable transient expression across different leaves of *N. tabacum* (Supplementary Fig. 2), consistent with the notion that AS109 is less sensitive to leaf physiological age (Lopez-Agudelo et al., 2025). This robustness is particularly important for assays with inherently variable readouts, such as immune receptor activation and pathogen infection, where minimizing leaf-to-leaf variation across biological and technical replicates is critical for reliable quantification.

**Fig. 1:**
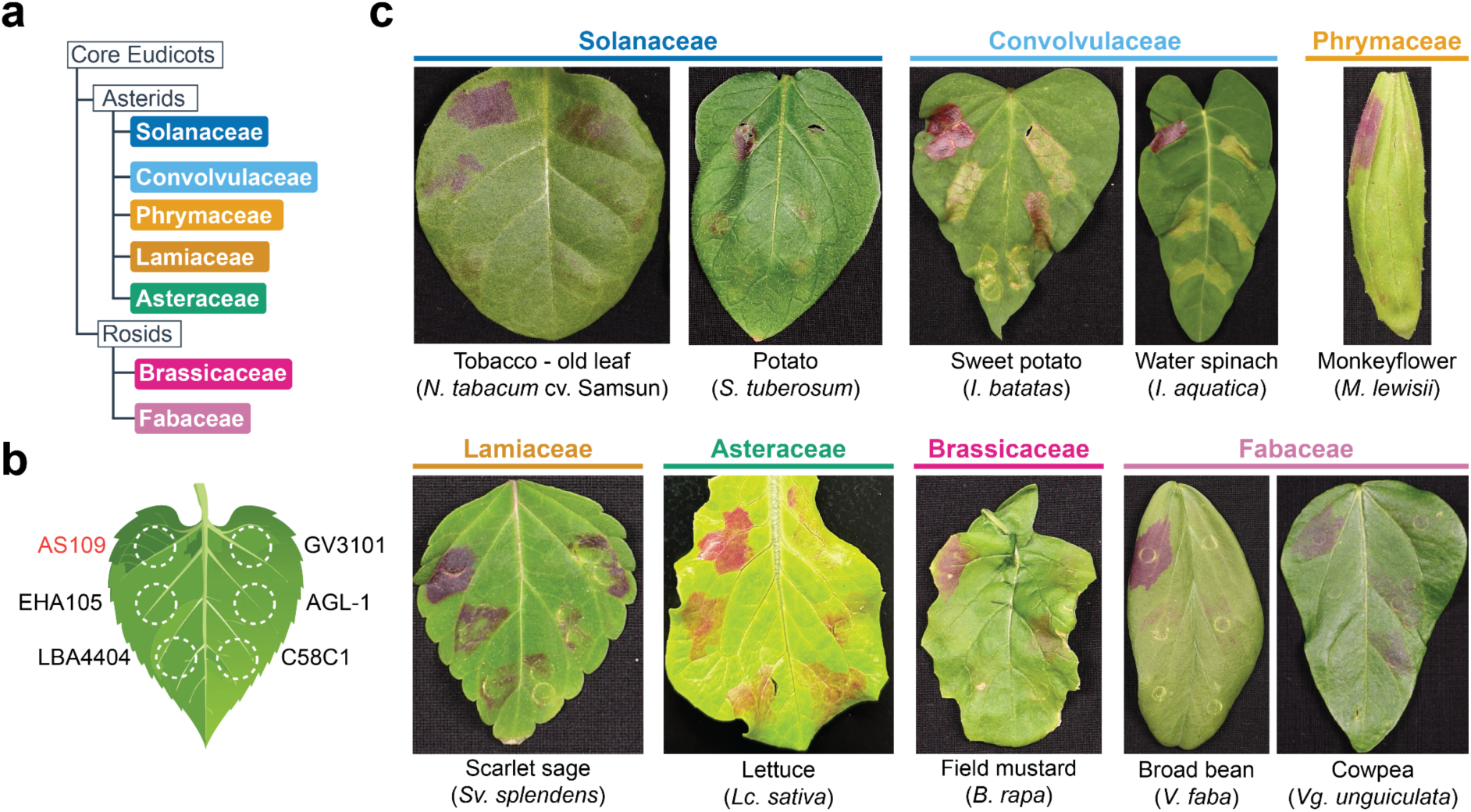
*Rhizobium rhizogenes* AS109 enables transient gene expression across diverse dicot species. **(a)** Simplified phylogenetic tree showing the evolutionary relationships of the plant families tested for RUBY transient expression. **(b)** Schematic representation of the infiltration layout used to compare six different agrobacterial strains. AS109 is positioned at the top left (highlighted in red). **(c)** Representative photographs of leaves showing transient expression of the RUBY reporter. Results display successful AS109-mediated expression across seven plant families: Solanaceae, Convolvulaceae, Phrymaceae, Lamiaceae, Asteraceae, Brassicaceae, and Fabaceae. All species tested and betalain quantification is shown in Supplementary Figs. 1-4.

We next examined the applicability of AS109 to several *Ipomoea* species in the family Convolvulaceae. While none of the other tested strains mediated clear and consistent transient reporter expression, AS109 enabled efficient transient expression in young expanding leaves of sweet potato (*I. batatas*), water spinach (*I. aquatica*), and *I. hederacea*, but not in *I. alba* (Fig. 1c and Supplementary Fig. 1d). Notably, AS109 induced fewer necrotic symptoms compared with other strains in *Ipomoea* spp., which may partly explain its higher efficiency in these species. This prompted us to extend our analysis to other asterid families and as well as to Caryophyllales (Supplementary Fig. 3a,b). AS109 mediated efficient transient expression in leaves of monkey flower (*Mimulus lewisii*; Phrymaceae), scarlet sage (*Salvia splendens*; Lamiaceae), sesame (*Sesamum indicum*; Pedaliaceae), and lettuce (*Lactuca sativa*; Asteraceae) (Fig. 1c and Supplementary Fig. 3c,d). In contrast, the other tested laboratory strains exhibited markedly lower expression efficiency in scarlet sage and lettuce, and showed little to no RUBY expression in monkey flower and sesame (Fig. 1c and Supplementary Fig. 3c,d). However, none of the tested laboratory strains, including AS109, mediated clear expression in snapdragon (*Antirrhinum majus*; Plantaginaceae), coffee (*Coffea canephora*; Rubiaceae), or red quinoa (*Chenopodium formosanum*; Amaranthaceae) (Supplementary Fig. 3e).

Next, we explored the potential application of AS109 in species from the Brassicaceae and Fabaceae, two rosid plant families that include many economically important crops (Fig. 1a and Supplementary Fig. 4a,b). Although AGL-1 and GV3101 mediated weak transient expression in field mustard (*Brassica rapa*; Brassicaceae), AS109 consistently produced stronger transient expression across different leaves (Fig. 1c and Supplementary Fig. 4c). We then tested eight species from the Fabaceae family. AS109 mediated consistent transient expression in alfalfa (*Medicago sativa*), fenugreek (*Trigonella foenum-graecum*), snow pea (*Pisum sativum*), common bean (*Phaseolus vulgaris*), faba bean (*Vicia faba*) and cowpea (*Vigna unguiculata*), but not soybean (*Glycine max*) or mung bean (*Vigna radiata*) (Fig. 1c and Supplementary Fig. 4d). In contrast, the other tested laboratory strains mediated low or inconsistent transient expression across these species (Supplementary Fig. 4d). Although AS109 did not support transient expression in every species tested, these results demonstrate that AS109 outperforms commonly used laboratory *Agrobacterium* strains in the majority of tested plants, underscoring its broad utility for transient expression across diverse plant taxa.

### AS109 facilitates functional assays in the non-model species faba bean and lettuce

To further explore whether AS109 can enable functional study of non-model species, we tested important transient assays in faba bean, owing to its amenability to transient expression in our initial screen and the global importance of Fabaceae crop species (Fig. 1c) (Jayakodi et al., 2023). Using the RUBY reporter, we verified that AS109 enables stronger transient expression in the first (most apical) and second leaf of faba bean compared to *A. fabrum* GV3101 and C58C1 (Fig. 2a,b). To test whether AS109 facilitates cell biology study in faba bean, we examined GFP localisation in the leaf epidermal cells by agroinfiltration of GV3101 and AS109 (Fig. 2c,d). Confocal microscopy revealed that AS109-mediated transient expression leads to stronger, more consistent GFP signals across faba bean epidermal cells. Importantly, AS109-mediated transient expression led to clear GFP signal in key subcellular compartments, the nucleus and cytoplasm (Fig. 2c). In addition to overexpression assays, we tested whether AS109-mediated expression could transiently knockdown genes in faba bean by RNAi, using previously established hairpin RNA constructs (Yan et al., 2012). To test RNAi in faba bean using AS109, we co-expressed GFP and mCherry fluorescent markers with hairpin GFP and GUS constructs (Fig. 2e,f). Co-expression with the hairpin GFP construct led to a significant reduction in GFP fluorescence compared to the hairpin GUS control, but had no significant effect on fluorescence from the mCherry control. We verified successful RNAi of GFP by RT-PCR, which showed a loss of GFP mRNA in leaves co-expressing hairpin GFP, but not with hairpin GUS (Fig. 2g). These results demonstrate that AS109 enables key transient assays in the non-model plant faba bean, facilitating the study of protein localization and genetics in an important Fabaceae crop species.

**Fig. 2:**
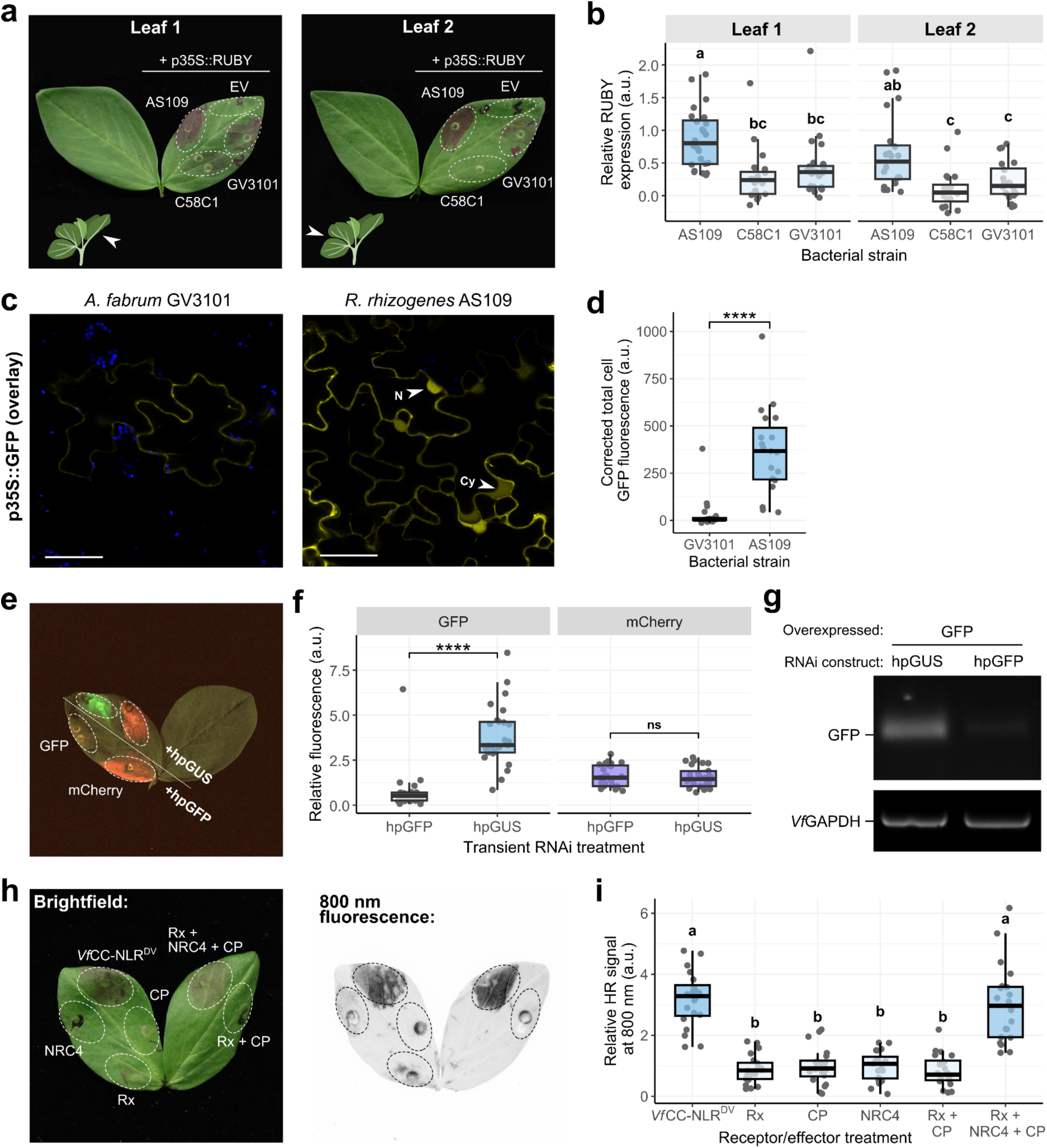
AS109 facilitates functional assays in the non-model species faba bean. **(a)** A representative image of the first and second faba bean leaf expressing RUBY by agroinfiltration of *A. fabrum* GV3101::pMP90, C58C1::pTiB6S3ΔT or *R. rhizogenes* AS109 compared to an empty vector (EV) control. **(b)** Box and dot plot depicting quantified betalain accumulation, normalised to the EV control, across three agroinfiltrations (n=28). Different letters above box plots represent statistically significant differences between treatments (p<0.05, one-way ANOVA with *post-hoc* Tukey HSD). **(c)** Representative confocal micrographs of faba bean epidermal cells expressing p35S::GFP by agroinfiltration of *A. fabrum* GV3101 or *R. rhizogenes* AS109. White arrows indicate GFP signal in the nucleus (N) and cytoplasm (Cy). Chloroplasts were visualised by autofluorescence at 648-709 nm emission. Scale bars represent 50 µm and both micrographs are single plane images. **(d)** Box and dot plot depicting corrected total cell GFP fluorescence from faba bean epidermal cells expressing p35S::GFP by infiltration of *A. fabrum* GV3101 or *R. rhizogenes* AS109 (n=20 cells). Asterisks above box plots represent statistically significant differences between treatments (p<0.05, Student’s t-test). **(e)** Representative stacked fluorescence image of an agroinfiltrated faba bean leaf at 532, 602 and 700 nm emission to visualise GFP fluorescence, mCherry fluorescence, and brightfield, respectively. **(f)** A box and dot plot depicting quantified GFP and mCherry fluorescence, normalised to background leaf fluorescence, at 532 and 602 nm emission, respectively. Agroinfiltrations were performed three times (n=25). **(g)** Validation of transient knockdown of GFP expression in faba bean by RT-PCR. RT-PCR was performed on cDNA generated from the total RNA of four infiltrated leaves. Glyceraldehyde 3-phosphate dehydrogenase (GAPDH) gene expression served as an internal control. **(h)** A representative image of a faba bean leaf under brightfield and under 800 nm emission, showing HR from expressing combinations of Rx, CP and *Nb*NRC4. Expression of an autoactive faba bean NLR (*Vf*CC-NLR^DV^) served as a positive control. **(i)** Box and dot plot showing the quantified HR signal at 800 nm emission from infiltrated faba bean leaves, normalised to the background leaf signal. Agroinfiltrations were performed three times (n=20).

To evaluate the generalisability of AS109-mediated transient assays, we performed the same assays in another non-model species, lettuce, a major leafy vegetable crop in the highly diverse Asteraceae family. Consistent with the results above, AS109 facilitated stronger transient expression in lettuce leaves than three commonly-used *A. fabrum* strains (Fig. 1c and Supplementary Fig. 5a,b). To check whether AS109 can enable cell biology studies in lettuce leaves, we compared GFP expression and localisation by agroinfiltration with AS109 and C58C1 (Supplementary Fig. 5c,d). Confocal microscopy demonstrated that AS109 caused stronger GFP expression in the epidermal cells than C58C1, evidenced by significantly higher total cell GFP fluorescence, and allows clearer visualisation of the nuclear compartment. To test AS109-mediated RNAi, we co-expressed GFP with hairpin GFP and GUS constructs in lettuce leaves (Supplementary Fig. 5e,f). Quantification of GFP fluorescence revealed a significant reduction upon co-expression with hairpin GFP but not hairpin GUS, suggesting that AS109 also allows transient gene knockdown in lettuce. RT-PCR of GFP mRNA also showed a reduction in GFP expression with hairpin GFP (Supplementary Fig. 5g). Taken together, AS109 also facilitates important transient assays in lettuce, extending the possibility of functional assays in multiple non-model crop species.

### AS109 enables rapid testing of immune receptors and elicitors in faba bean

Conventional *Agrobacterium*-mediated transient expression in *N. benthamiana* has made major contributions to the functional analysis of plant disease-resistance proteins, particularly nucleotide-binding leucine-rich repeat receptors (NLRs) and cell-surface pattern recognition receptors (PRRs), by enabling rapid tests of receptor autoactivity, signaling requirements, and effector recognition (Bally et al., 2018). To evaluate the use of AS109-mediated transient expression for studying immunity in non-model species, we tested the function of autoactive NLRs in faba bean leaves. AS109-mediated transient expression of autoactive solanaceous helper NLRs such as NRC2, NRC3, and NRC4 variants carrying mutations in the MHD motif, but not their WT variants, triggered cell death indicating that NRC helpers can recapitulate their autoactive functions in Fabaceae (Supplementary Fig. 6a,b) (Wu et al., 2017). Consistently, expression of an autoactive, but not WT, endogenous coiled-coil (CC)-NLR variant cloned from faba bean (*Vf*CC-NLR) triggered cell death. Expression of autoactive NRC4 with additional L to E mutations in the CC domain (*Nb*NRC4^DV/3E^) did not cause HR, showing that NRC4 auto-activity in faba bean requires an intact CC domain, consistent with the earlier results obtained using *N. benthamiana* (Supplementary Fig. 6c,d) (Adachi et al., 2019). Using the non-cell-death-inducing NRC4 L9E mutant (*Nb*NRC4^DV/L9E^), we were able to monitor activated NLR resistosome clusters, observed as membrane-associated puncta, in contrast to the diffuse cytoplasmic distribution of resting, unactivated NRC4 (Supplementary Fig. 6e).

Solanaceae NLRs are relatively well studied compared with those from most other plant families and represent a potential source of novel disease-resistance genes for non-solanaceous crops (Du et al., 2025; Oh et al., 2024). However, divergence in required signaling pathways can render many transferred NLRs non-functional, a phenomenon known as restricted taxonomic functionality (Guo and Jones, 2026). To assess whether AS109-mediated transient expression enables rapid evaluation of NLR transferability, and to distinguish NLRs that require additional components from those that do not, we performed proof-of-concept experiments by expressing two solanaceous NLRs together with their corresponding avirulence factors in faba bean leaves.

We first tested the cell death function of the singleton NLR Rpi-vnt1 in faba bean leaves (Supplementary Fig. 7). Rpi-vnt1 was originally cloned from *Solanum venturii* and provides resistance to *Phytophthora infestans* by recognition of the effector Avr-vnt1 (Foster et al., 2009). Transient expression of Rpi-vnt1 under its native promoter in faba bean led to cell death upon co-expression with its cognate effector Avr-vnt1, showing that effector recognition of this singleton NLR can be directly transferred from Solanaceae to Fabaceae without additional components.

Next, we tested the cell death function of the sensor NLR Rx, which recognises the PVX coat protein (CP) and requires NRC helper NLRs to trigger cell death (Fig. 2h,i) (Wu et al., 2017; Contreras et al., 2024). Transient expression of Rx with CP in faba bean triggered cell death only when *Nb*NRC4 was co-expressed, showing that the cell death specificity of sensor NLRs can also be transferred by providing the required helper NLR. Together, these results demonstrate that both singleton NLRs and sensor-helper NLR pairs from solanaceous plants can be moved into faba bean to provide their native effector recognition and cell death functions. This establishes Solanaceae as a potential source for novel disease-resistance genes for other plant families and highlights AS109 as a rapid platform for testing restricted taxonomic functionality across divergent plant families.

We then tested whether AS109 can be used to assess cell-surface recognition of apoplastic pathogen elicitors and effectors in faba bean. We screened a panel of 23 secreted proteins from oomycete, fungal, bacterial, and nematode plant pathogens, which are required for pathogen virulence and have been reported to trigger immune responses in *N. benthamiana* (Supplementary Table 2). Transient expression of these elicitors triggered varying degrees of cell death in faba bean leaves, including strong cell death caused by the cytotoxic protein *Zt*NLP (Supplementary Fig. 8) (Motteram et al., 2009). Remarkably, the conserved effector XEG1 from *Phytophthora sojae* triggered weak cell death in faba bean, suggesting that this species may harbour an unknown PRR that recognises XEG1. These data show that AS109 also enables testing of cell-surface recognition in faba bean, making this a powerful platform to screen for novel sources of disease resistance in legumes.

### AS109-mediated expression reveals a novel host-pathogen interface in faba bean

Finally, we asked whether AS109-mediated transient expression in faba bean could support infection cell biology assays, an application that has been particularly powerful in *N. benthamiana* for revealing novel defence and immune-suppression mechanisms at the host-pathogen interface (Bozkurt and Kamoun, 2020). To test this, we transiently expressed the potato REMORIN marker *St*REM1.3 fused to RFP in faba bean leaves and subsequently infected detached leaves with a transgenic *Phytophthora palmivora* line expressing ER-targeted YFP (Rey et al., 2014). Imaging at 1-2 days post-inoculation revealed clear accumulation of *St*REM1.3 around *P. palmivora* haustoria, marking the extrahaustorial membrane, the specialised host-derived membrane interface that surrounds invading oomycete haustoria (Supplementary Fig. 9) (Bozkurt et al., 2014). These results show that AS109-mediated transient expression in faba bean is compatible not only with immune receptor assays, but also with infection cell biology approaches for visualising host-pathogen interfaces during infection.

## Conclusions

In summary, we demonstrate that the *R. rhizogenes* strain AS109 enables robust transient expression across a substantially broader spectrum of wild and crop dicotyledonous species than commonly used laboratory *Agrobacterium* strains. By overcoming long-standing host-range limitations, AS109 provides a versatile and accessible platform for molecular and cell biological analyses in non-model plant species that have remained largely recalcitrant to transient transformation. Its performance across diverse phylogenetic lineages creates new opportunities for comparative studies of plant immunity, signaling, and cell biology in an evolutionary context.

Beyond basic research, AS109 offers a practical route for rapid gene validation, trait screening, and pathway engineering in crops and wild relatives. In particular, functional interrogation of natural diversity in wild species may accelerate the identification of candidate resistance genes for incorporation into pre-breeding and crop improvement pipelines (Liu et al., 2025; Lin et al., 2023). The success of *Agrobacterium*-mediated transient assays in *N. benthamiana* has helped make Solanaceae a particularly productive system for the discovery and mechanistic analysis of disease-resistance genes, with especially strong progress in R-gene identification and cloning in crops such as potato (Jones et al., 2024; Zhu et al., 2012). By extending comparable rapid assays into Fabaceae, AS109 could similarly accelerate the discovery and functional testing of legume disease-resistance traits and mechanisms. As protein structure prediction and comparative genomics continue to accelerate hypothesis generation in plant science, AS109 provides a much-needed platform for rapid functional validation across diverse species, helping to bridge the gap between computational prediction, mechanistic insight, and translational crop improvement.

AS109 also helps overcome redundancy-related challenges in plant immune assays. By enabling immune receptors to be tested across divergent crop families, it offers a tractable heterologous system for dissecting receptor function in genetic backgrounds with reduced endogenous redundancy. This could be especially powerful for resolving helper-sensor NLR relationships, which in native hosts often require complex knockdown or knockout strategies to separate overlapping helper activities (Adachi and Kamoun, 2022). Collectively, our work establishes AS109 as a broadly applicable tool that expands the experimental repertoire of plant biotechnology and accelerates the discovery of novel disease-resistance traits and mechanisms for crop improvement.

## Acknowledgements

We thank Dr. Chun-Neng Wang (Institute of Ecology and Evolutionary Biology, National Taiwan University) for sharing the seeds of *Mimulus lewisii*, Dr. Heng-Ming Ting (Institute of Plant Biology, National Taiwan University) for sharing the seeds of *V. radiata*, Dr. Na-Sheng Lin (Institute of Plant and Microbial Biology, Academia Sinica) for sharing the seeds of *Chenopodium formosanum*, Dr. Ya-Chun Chang (Department of Plant Pathology and Microbiology, National Taiwan University) for sharing the seeds of *Nicotiana tabacum*.

We thank the Imperial College FILM facility for their technical expertise and provision of our microscopy equipment.

## Funding

Biotechnology and Biological Sciences Research Council (BBSRC) BB/X016382/1 (T.O.B.)

Bezos Earth Fund through the Bezos Centre for Sustainable Protein BCSP/IC/001 (T.O.B.)

Academia Sinica Grand Challenge Program AS-GC-111-L02 (E.M.L., C.H.K. and C.H.W.)

Intramural fund of the Institute of Plant and Microbial Biology, Academia Sinica (E.M.L., C.H.K. and C.H.W.)

The Niab Trust, through an independent fellowship (L.D.)

Royal Society University Research Fellowship URF\R1\251226 (J.K.).

European Research Council (ERC) Starting Grant 101220795 – PREDESIGNX (J.K.)

## Author contributions

Conceptualization: F.K., J.C.L.A., C.S., L.D., C.H.W., T.O.B.

Methodology: F.K., J.C.L.A., C.S., M.H.A., T.I., A.C., N.E., S.J., C.V., Y.N.S., W.J.L., S.B., A.T.

Investigation: F.K., J.C.L.A., C.S., M.H.A., T.I., E.L.H.Y., A.C., N.E., S.J., C.V., Y.N.S., W.J.L.

Visualization: F.K., J.C.L.A., T.I., E.L.H.Y.

Funding acquisition: C.H.K., E.M.L., J.K., L.D., C.H.W., T.O.B.

Project administration: F.K., C.H.K., E.M.L., J.K., L.D., C.H.W., T.O.B.

Supervision: F.K., J.C.L.A., C.H.K., E.M.L., J.K., L.D., C.H.W., T.O.B.

Writing – original draft: F.K., J.C.L.A., C.S., L.D., C.H.W., T.O.B.

Writing – review & editing: F.K., J.C.L.A., C.S., C.H.K., E.M.L., J.K., L.D., C.H.W., T.O.B.

## Data and materials availability

Constructs generated for this study will be subjected to material transfer agreements (MTAs) and made available upon request. All data are available in the main text or the supplementary materials.

## Supplementary figures

**Supplementary Fig. 1:**
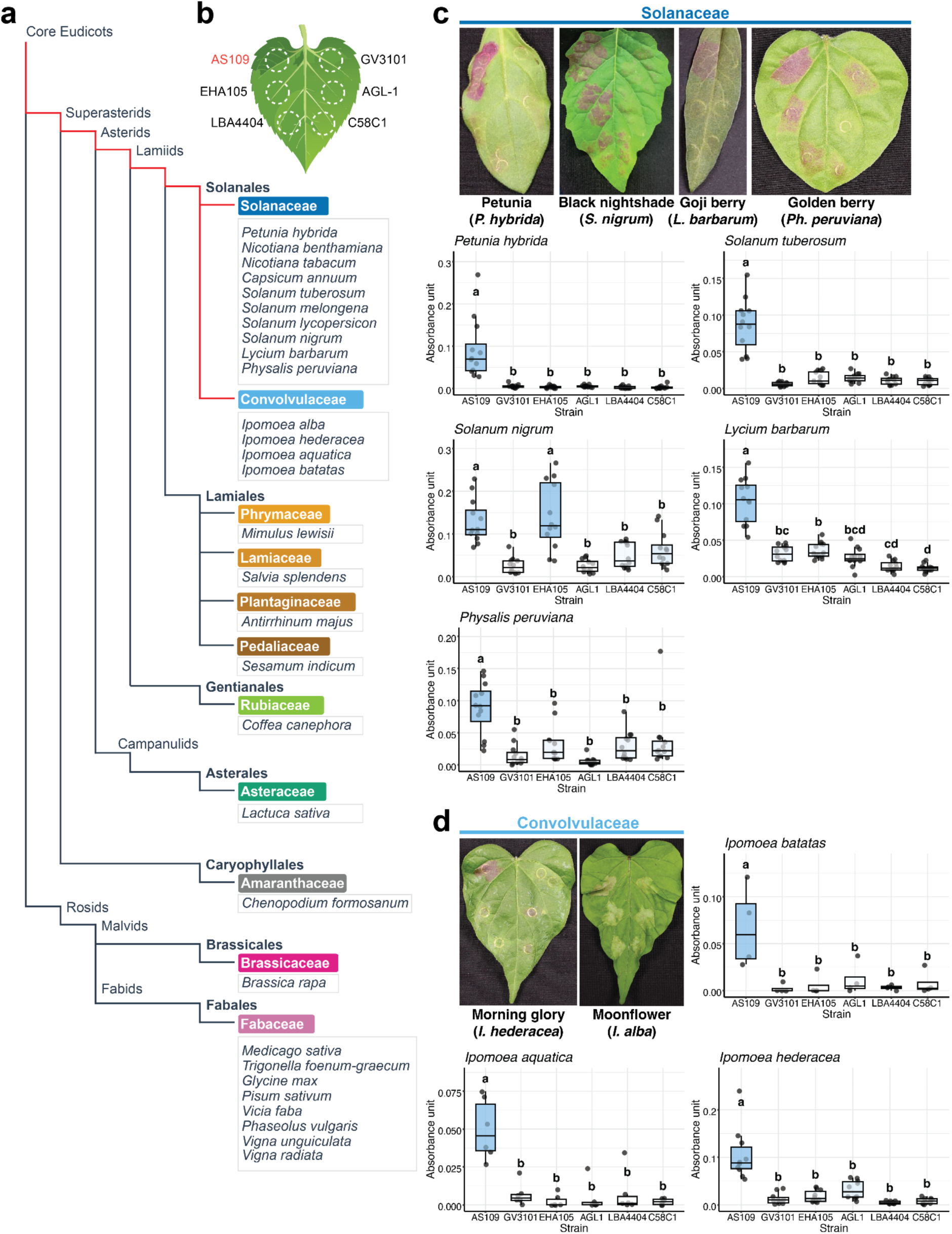
AS109 mediates successful transient gene expression within the Solanales. **(a)** Detailed phylogeny of the 30 angiosperm species evaluated for RUBY transient expression. The selection spans the Core Eudicots, including Superasterids (Caryophyllales and Asterids) and Rosids. The Solanales lineage (Solanaceae and Convolvulaceae) is highlighted in red. **(b)** Schematic representation of the infiltration layout used to compare six different agrobacterial strains. AS109 is positioned at the top left (highlighted in red). **(c, d)** Representative photographs and betalain quantification for Solanaceae **(c)** and Convolvulaceae **(d)**. Photographs display species that were not shown in Figure 1. Box plots represent the total betalain absorbance measurement in each infiltration spot (Solanaceae n=12; *I. batatas* n=4; *I. aquatica* n=6; *I. hederacea* n=10). Different letters above box plots represent statistically significant differences between strains (p<0.05, one-way ANOVA with *post-hoc* Tukey HSD).

**Supplementary Fig. 2:**
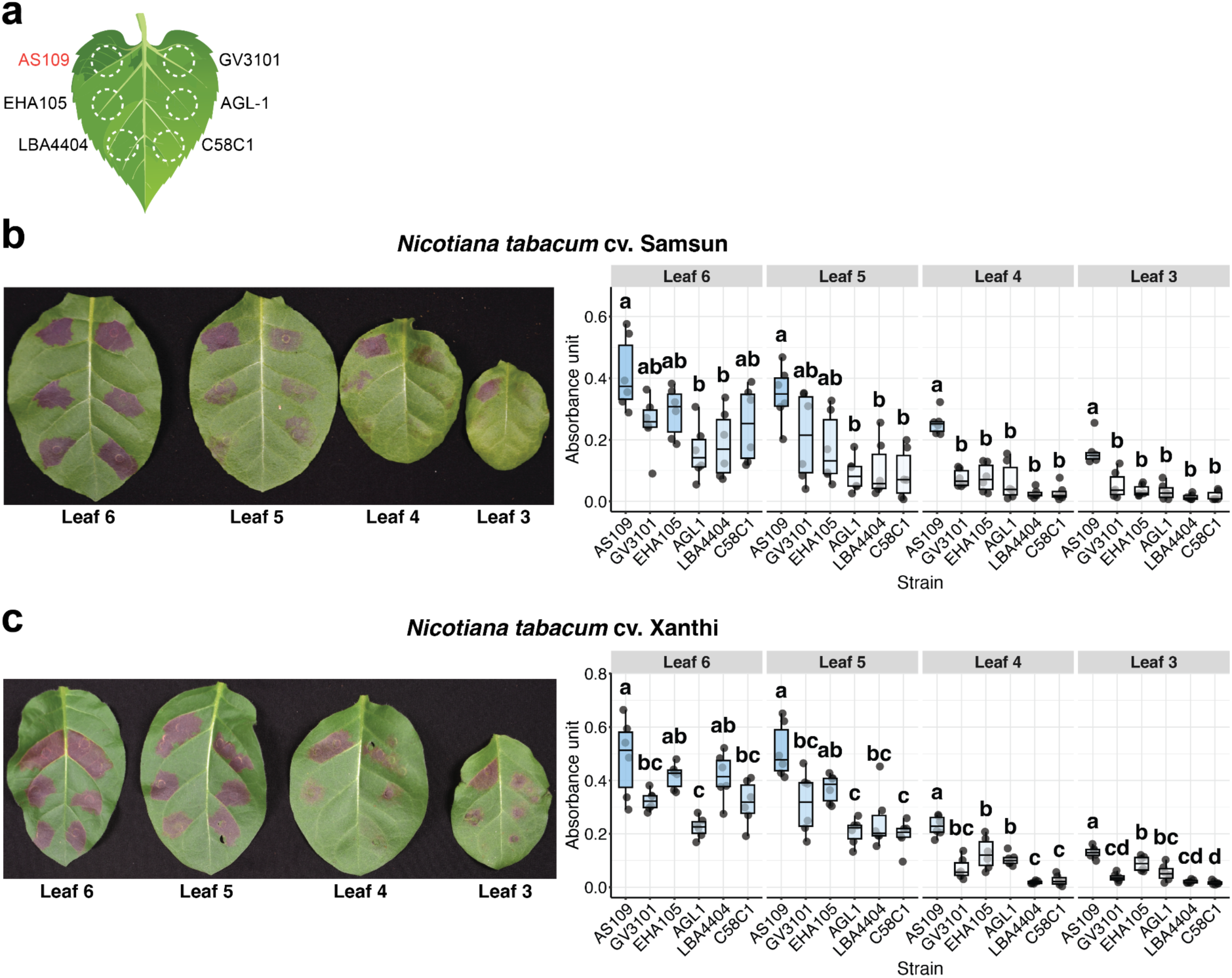
AS109 mediates reliable transient expression across different *N. tabacum* leaves. **(a)** Schematic representation of the infiltration layout used to compare six different agrobacterial strains. AS109 is positioned at the top left (highlighted in red). **(b, c)** Representative photographs and betalain quantification for leaves of different physiological ages in *Nicotiana tabacum* cv. Samsun **(b)** and cv. Xanthi **(c)**. Box plots represent the total betalain absorbance measurement in each infiltration spot from leaves 3 (older) through 6 (younger) (n=6). Different letters above box plots represent statistically significant differences between strains (p<0.05, one-way ANOVA with *post-hoc* Tukey HSD).

**Supplementary Fig. 3:**
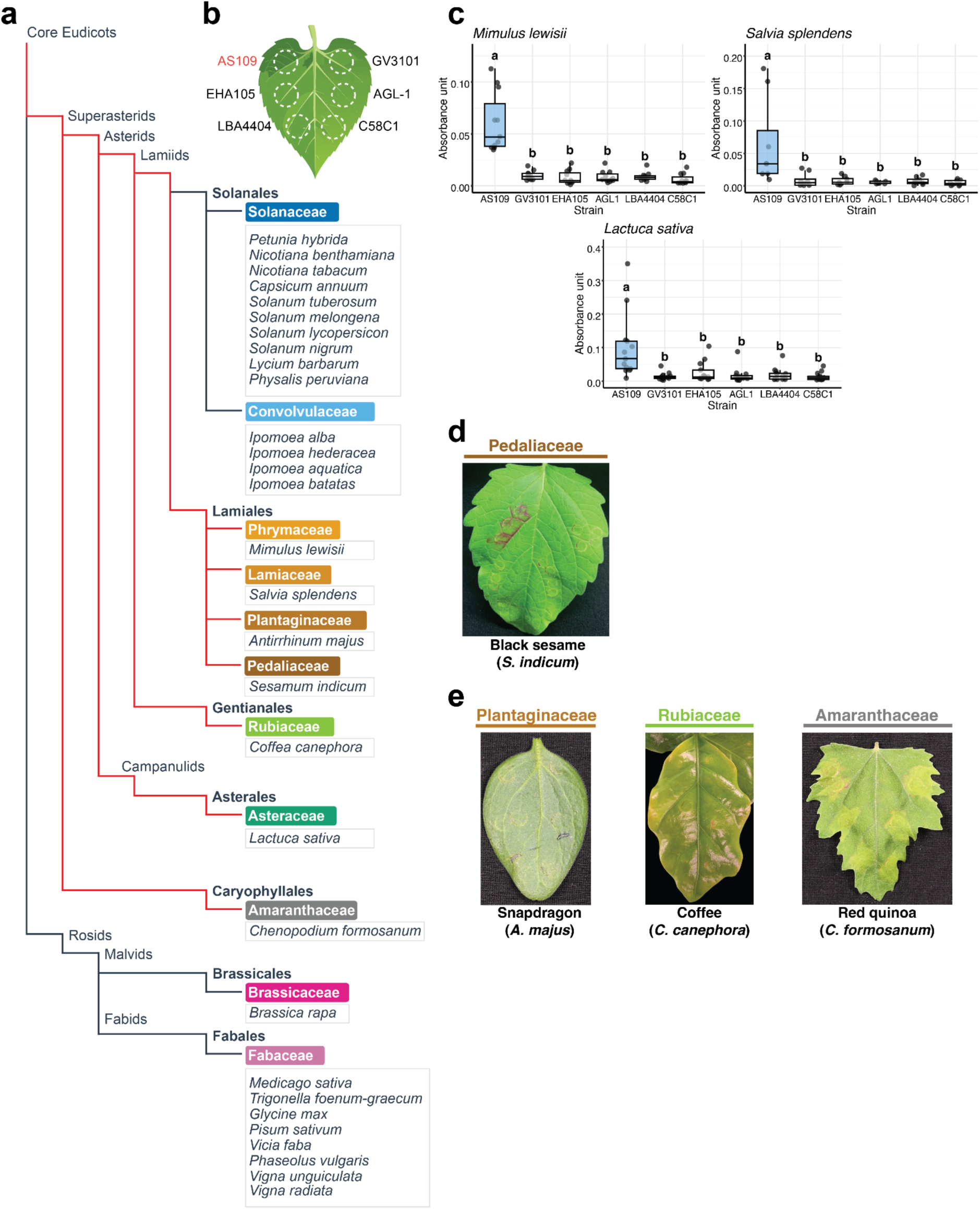
AS109 mediates successful transient gene expression across several non-Solanales Superasterid families. **(a)** Detailed phylogeny of the 30 angiosperm species evaluated for RUBY transient expression. The selection spans the Core Eudicots, including Superasterids (Caryophyllales and Asterids) and Rosids. The Superasterids lineages detailed in this figure (Lamiales, Gentianales, Asterales and Caryophyllales) are highlighted in red. **(b)** Schematic representation of the infiltration layout used to compare six different agrobacterial strains. AS109 is positioned at the top left (highlighted in red). **(c)** Box plots representing the total betalain absorbance measurement in each infiltration spot in species with successful RUBY expression (*M. lewisii* n=11; *Sv. splendens* n=8; *Lc. sativa* n=13). Different letters above box plots represent statistically significant differences between strains (p<0.05, one-way ANOVA with *post-hoc* Tukey HSD). **(d, e)** Representative photographs of species exhibiting weak **(d)** or undetectable **(e)** RUBY expression.

**Supplementary Fig. 4:**
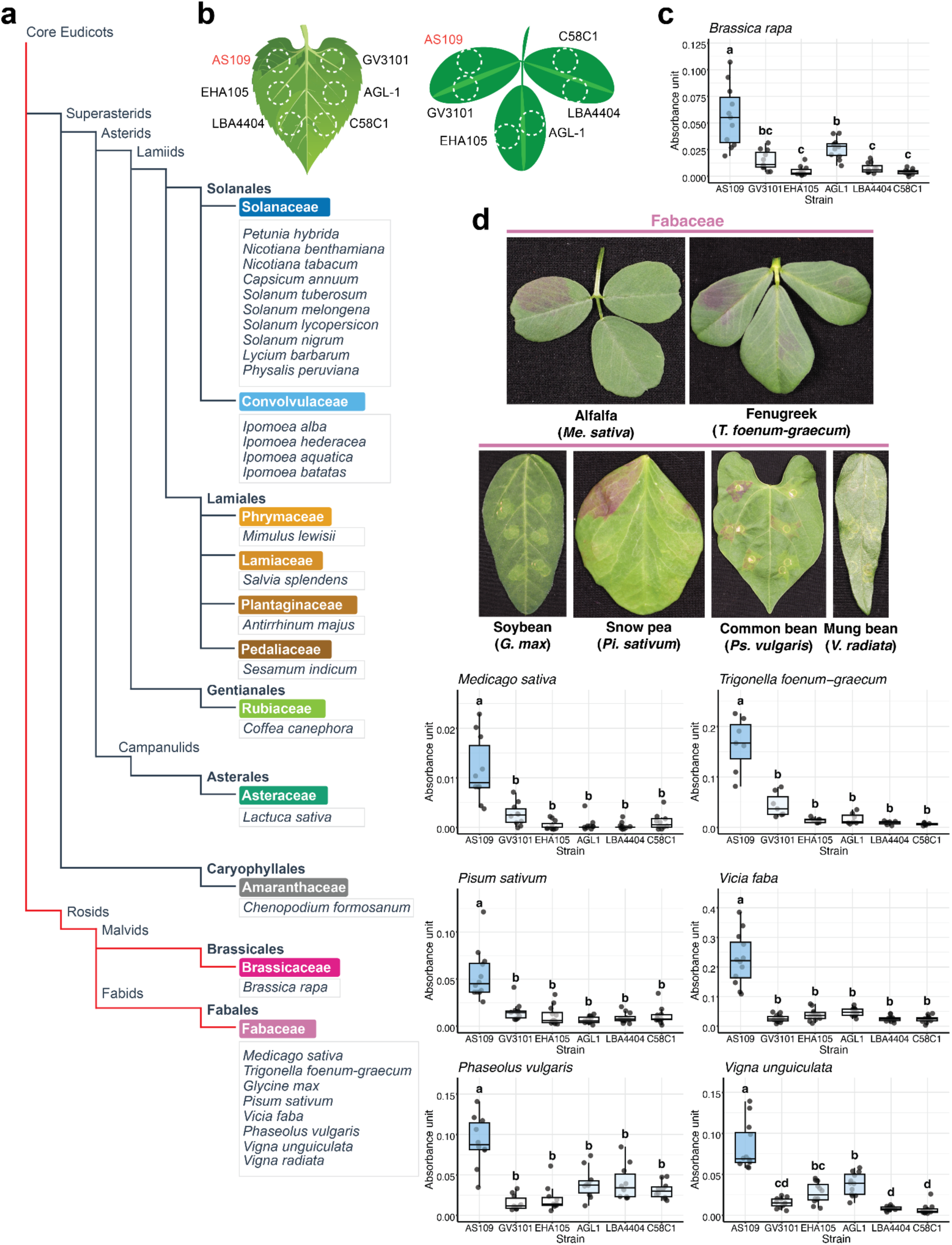
AS109 mediates successful transient gene expression across Brassicaceae and Fabaceae. **(a)** Detailed phylogeny of the 30 angiosperm species evaluated for RUBY transient expression. The selection spans the Core Eudicots, including Superasterids (Caryophyllales and Asterids) and Rosids. The Rosids lineage (Brassicaceae and Fabaceae) is highlighted in red. **(b)** Schematic representation of the infiltration layout used to compare six different agrobacterial strains. AS109 is positioned at the top left (highlighted in red). **(c, d)** Representative photographs and betalain quantification for Brassicaceae **(c)** and Fabaceae **(d)**. Photographs display species that were not shown in Figure 1. Box plots represent the total betalain absorbance measurement in each infiltration spot (*B. rapa* n=11; *Me. sativa* n=10; *T. foenum-graecum* n=7; *Pi. sativum*, *V. faba* and *Vg. unguiculata* n=12; *Ps. vulgaris* n=10). Different letters above box plots represent statistically significant differences between strains (p<0.05, one-way ANOVA with *post-hoc* Tukey HSD).

**Supplementary Fig. 5:**
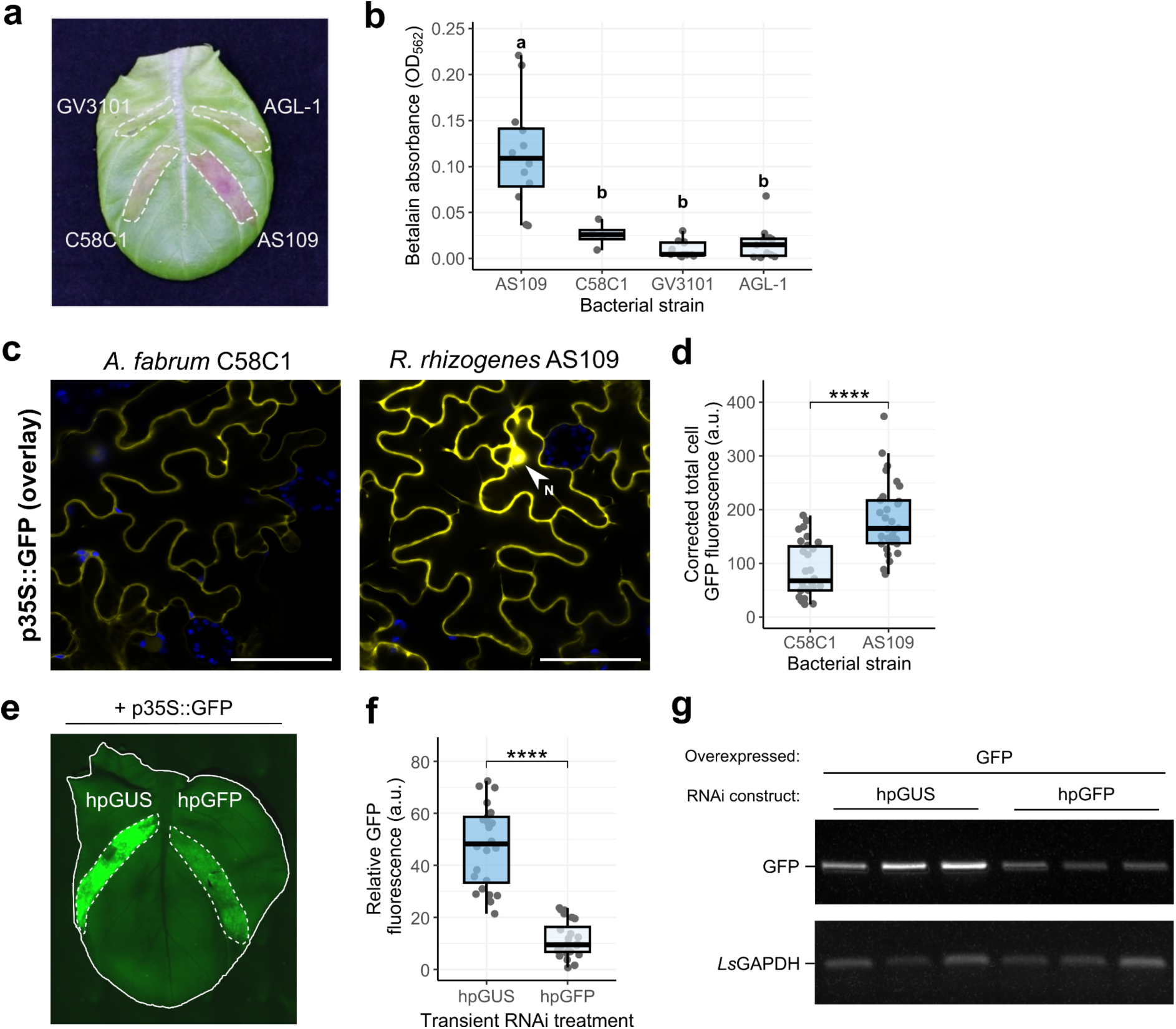
AS109 facilitates functional assays in the non-model species lettuce. **(a)** Representative image of a lettuce leaf expressing RUBY by agroinfiltration with the indicated *A. fabrum* strains and *R. rhizogenes* AS109. **(b)** Box and dot plot depicting quantified betalain concentration measured by OD_562_ nm across two agroinfiltrations (n=12). Different letters above box plots represent statistically significant differences between treatments (p<0.05, one-way ANOVA with *post-hoc* Tukey HSD). **(c)** Representative confocal micrographs of lettuce epidermal cells expressing p35S::GFP by agroinfiltration of *A. fabrum* C58C1::pTiB6S3ΔT or *R. rhizogenes* AS109. The white arrow indicates GFP signal in the nucleus (N). Chloroplasts were visualised by autofluorescence at 648-709 nm emission. Scale bars represent 50 µm and both micrographs are single plane images. **(d)** Box and dot plot depicting corrected total cell GFP fluorescence from lettuce epidermal cells expressing p35S::GFP by infiltration of *A. fabrum* C58C1 or *R. rhizogenes* AS109 (n=60 cells). Agroinfiltrations were repeated three times. Asterisks above box plots represent statistically significant differences between treatments (p<0.05, Student’s t-test with Bonferroni correction). **(e)** Representative GFP fluorescence (500-550 nm emission) image of an agroinfiltrated lettuce leaf expressing p35S::GFP with a hairpin GFP RNAi construct or a hairpin GUS control. **(f)** A box and dot plot depicting quantified GFP fluorescence, normalised to background leaf fluorescence at 532 emission. Agroinfiltrations were repeated twice (n=24). **(g)** Validation of transient knockdown of GFP expression in lettuce by RT-PCR. RT-PCR was performed on cDNA generated from the total RNA of three infiltrated leaves. Glyceraldehyde 3-phosphate dehydrogenase (GAPDH) gene expression served as an internal control.

**Supplementary Fig. 6:**
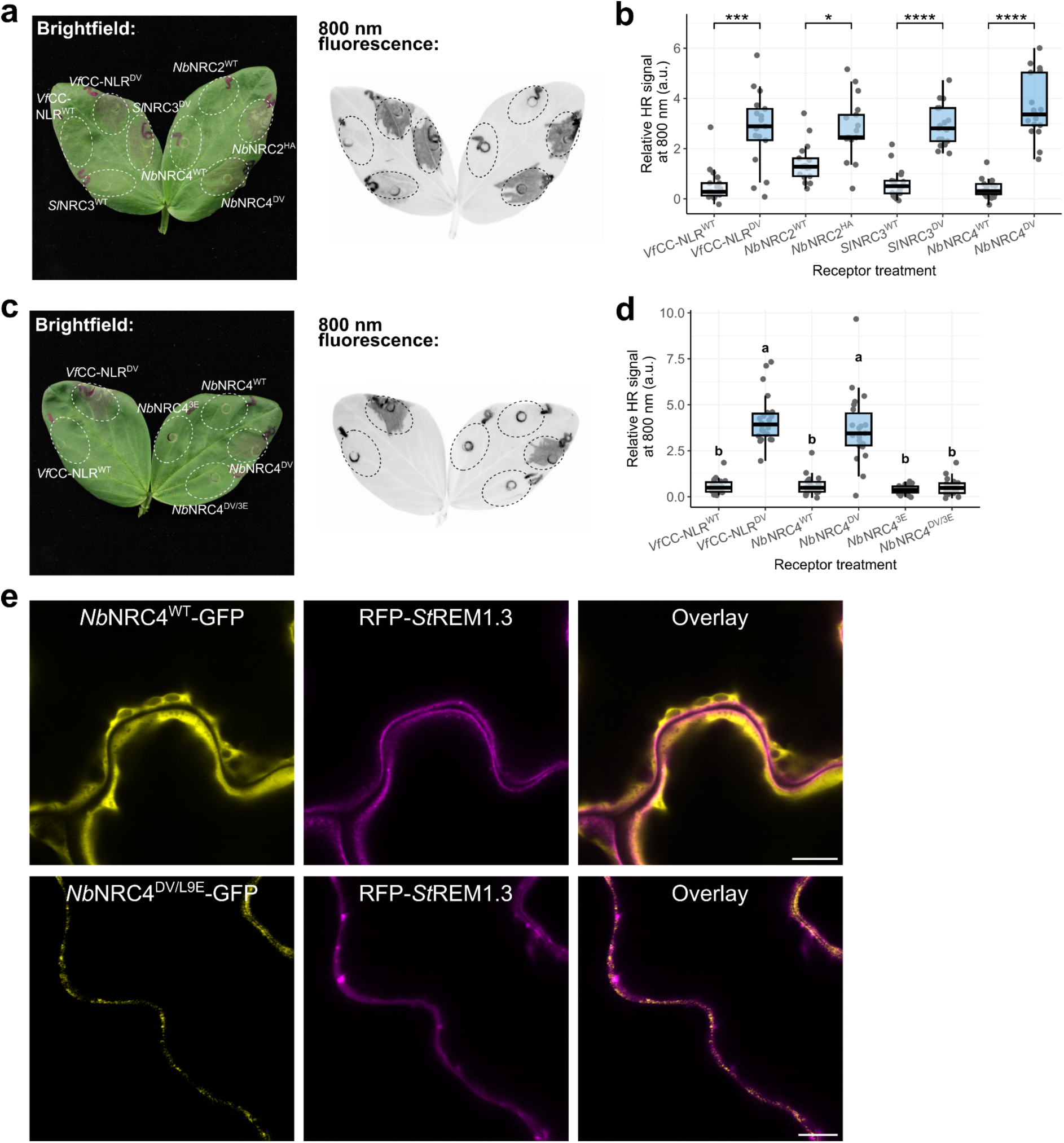
autoactive helper NLRs cause cell death and form puncta at the plasma membrane in faba bean. **(a)** Representative image of a faba bean leaf showing HR from expressing WT and autoactive helper NLRs under brightfield (left) and under 800 nm emission (right). Expression of the WT and autoactive variant of a faba bean NLR (*Vf*CC-NLR) served as a negative and positive control, respectively. **(b)** Box and dot plot showing the quantified HR signal from expressing WT and autoactive helper NLRs at 800 nm emission from infiltrated faba bean leaves, normalised to the background leaf signal. Asterisks above box plots represent statistically significant differences between treatments (p<0.05, Student’s t-test with Bonferroni correction). Agroinfiltrations were performed three times (n=17). **(c)** Representative image of a faba bean leaf showing HR from expressing *Nb*NRC4 mutants. Expression of the WT and autoactive variant of a faba bean NLR (*Vf*CC-NLR) served as a negative and positive control, respectively. **(d)** Box and dot plot showing the quantified HR signal of expressing *Nb*NRC4 mutants under 800 nm emission from infiltrated faba bean leaves, normalised to the background leaf signal. Different letters above box plots represent statistically significant differences between treatments (p<0.05, one-way ANOVA with *post-hoc* Tukey HSD). Agroinfiltrations were performed three times (n=22). **(e)** Representative confocal micrographs of faba bean epidermal cells transiently expressing *Nb*NRC4^WT^-GFP (n=5 images), or *Nb*NRC4^DV/L9E^-GFP (n=11 images) with RFP-*St*REM1.3. Scale bars represent 10 µm.

**Supplementary Fig. 7:**
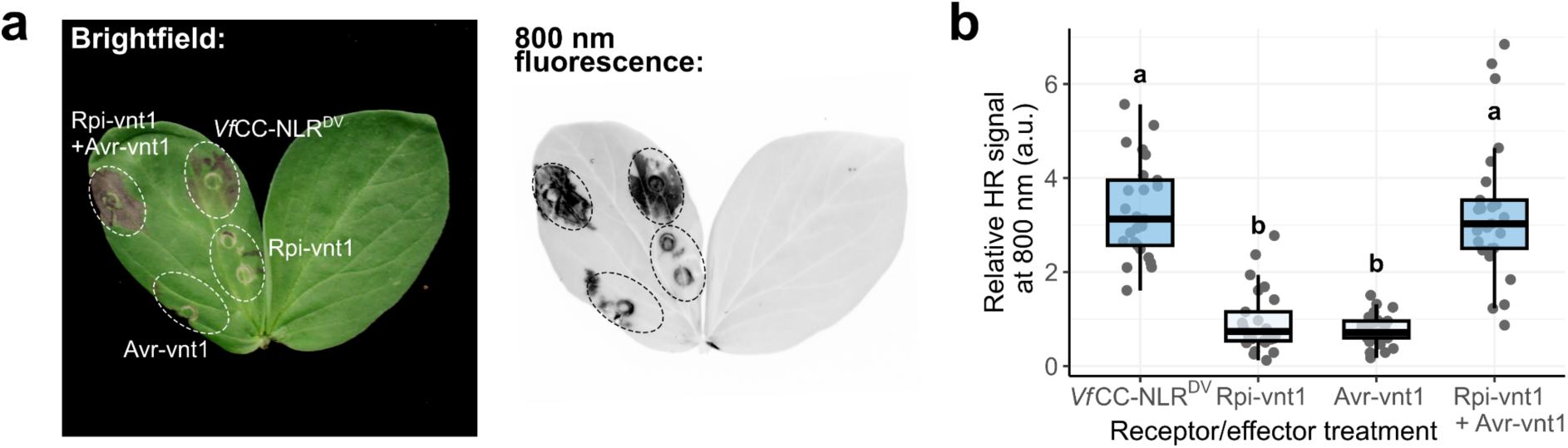
A singleton NLR from Solanaceae, Rpi-vnt1, performs its cell death function in faba bean. **(a)** Representative image of a faba bean leaf showing HR from expressing combinations of Rpi-vnt1 and Avr-vnt1 under brightfield (left) and under 800 nm emission (right). Expression of an autoactive faba bean NLR (*Vf*CC-NLR^DV^) served as a positive control. **(b)** Box and dot plot showing the quantified HR under 800 nm emission from infiltrated faba bean leaves, normalised to the background leaf signal. Different letters above box plots represent statistically significant differences between treatments (p<0.05, one-way ANOVA with *post-hoc* Tukey HSD). Agroinfiltrations were performed three times (n=25).

**Supplementary Fig. 8:**
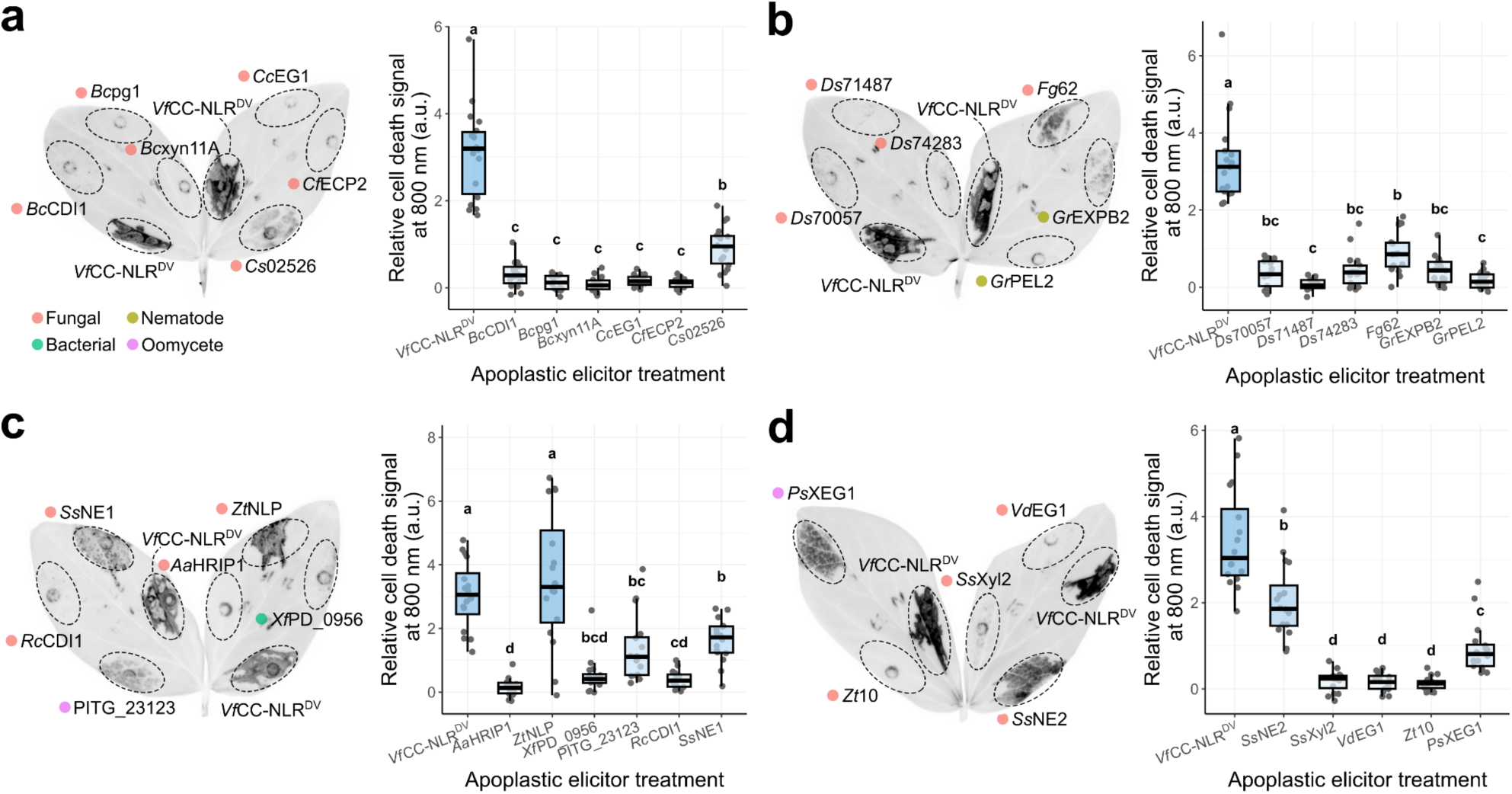
Apoplastic elicitors trigger cell death in faba bean. **(a-d)** Representative image of a faba bean leaf showing cell death from expressing apoplastic elicitors under 800 nm emission (left) and box and dot plot (right) showing the quantified cell death signal, normalised to the background leaf signal, for four groups of elicitors. Expression of the autoactive variant of a faba bean NLR (*Vf*CC-NLR) served as a positive control in each case. Agroinfiltrations were performed twice (n=16). Different letters above box plots represent statistically significant differences between treatments (p<0.05, one-way ANOVA with *post-hoc* Tukey HSD). Details of the apoplastic elicitors can be found in Supplementary Table 2.

**Supplementary Fig. 9:**
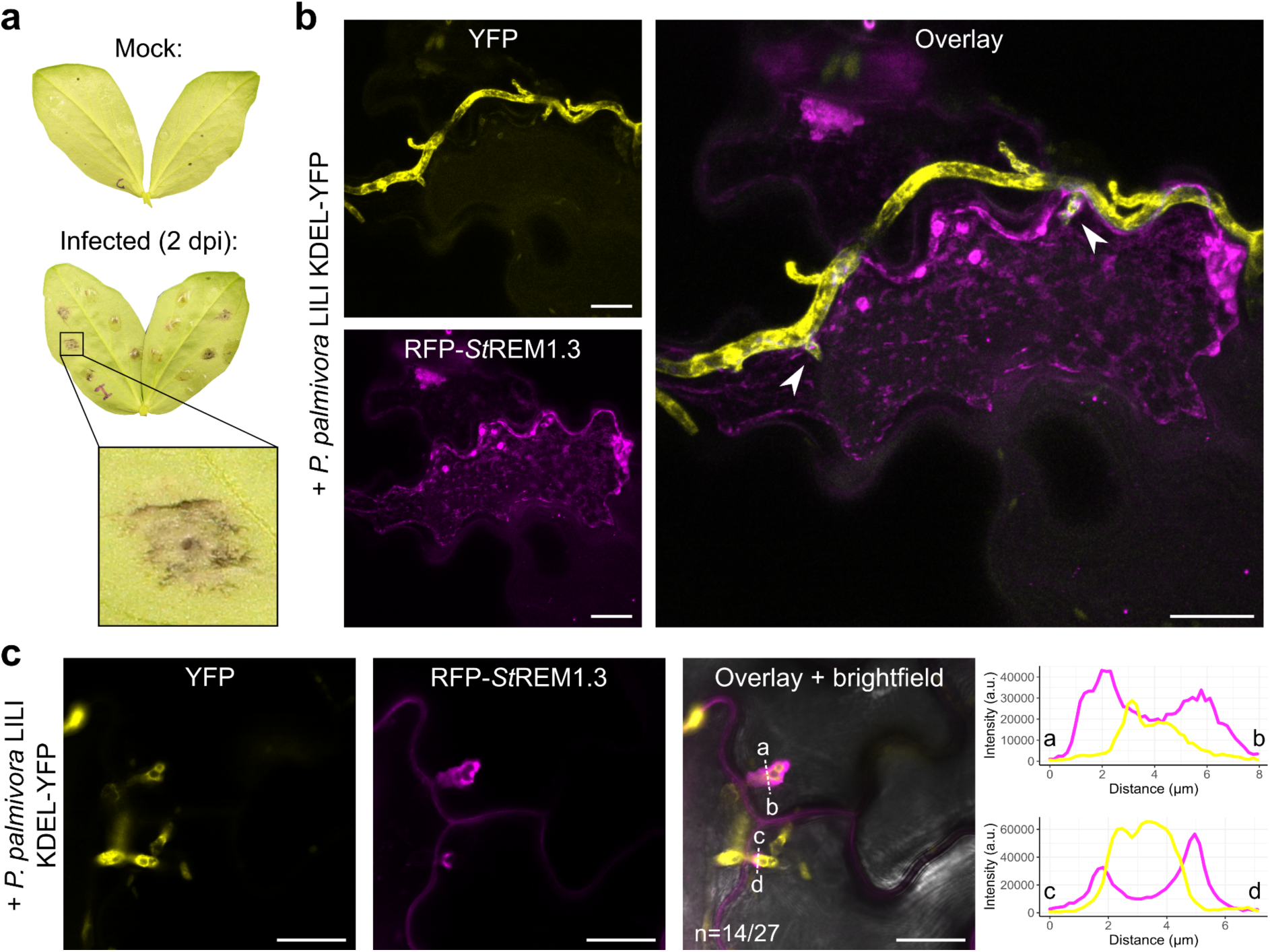
AS109-mediated expression reveals a novel host-pathogen interface in faba bean. **(a)** Representative image of faba bean leaves mock treated (upper) and infected with *Phytophthora palmivora* LILI KDEL-YFP spores (lower). Photographs were taken at 2 days post-inoculation (dpi). **(b)** Confocal micrograph showing an agroinfiltrated faba bean epidermal cell expressing RFP-*St*REM1.3 whilst infected by *P. palmivora*. The image shown is a maximum projection of z-stacks. Arrows indicate intracellular haustoria. Scale bars represent 20 µm. **(c)** Representative confocal micrographs of faba bean epidermal cells expressing RFP-*St*REM1.3 surrounding *P. palmivora* haustoria (14/27 haustoria, n=10 images). Transects correspond to the line intensity plots of relative fluorescence intensity across the marked distances. Scale bars represent 20 µm.

## Methods

### Plant growth conditions

All plant species were grown in climate-controlled growth chambers at 22-28°C with 16/8 hour day/night light cycles. Most plants were grown on a combination of soil and vermiculite mixed at a 3:1 ratio and were agroinfiltrated at 3-5 weeks old. Faba bean (cultivar ‘de Monica’) and lettuce (cultivar ‘Amica’) plants were grown in Levington’s F2 soil with sand and Sinclair’s 2-5 mm vermiculite, mixed in a 3:1 ratio. Faba bean plants were agroinfiltrated at 1-2 weeks old and lettuce were agroinfiltrated at 2-3 weeks old. The cultivars and origin of all plant material used in this study is detailed in Supplementary Table 1.

### Molecular cloning

Constructs made in this study were cloned using Golden Gate Modular Cloning (MoClo) with the MoClo Plant Parts (Engler et al., 2014). Level 0 gene inserts were produced by commercial gene synthesis (Twist Bioscience) except for *Vf*CC-NLR (gene Vfaba.Hedin2.R1.2g009560.1; Jayakodi et al., 2023), which was cloned by PCR from faba bean cDNA (Supplementary Table 3). Constructs were verified by sequencing before transformation into *A. fabrum* strains or *R. rhizogenes* AS109 by electroporation. Construct sequences were designed and verified in Geneious Prime (v2026.0.2). The remaining constructs were provided from previous studies (Supplementary Table 4).

### Agroinfiltration for transient expression

Agroinfiltration of all plant species was performed as previously described for *Nicotiana benthamiana* (Ibrahim et al., 2024). *Agrobacterium* strains containing binary plasmids were grown on LB plates containing selective antibiotics (100 μg/mL kanamycin or 100 μg/mL spectinomycin) and a secondary antibiotic to maintain the virulence plasmid according to the strain (Supplementary Table 5). *R. rhizogenes* AS109 containing binary plasmids was grown on 523 media plates containing selective antibiotics (100 μg/mL kanamycin or 200 μg/mL spectinomycin) and rifampicin (50 μg/mL). Bacterial cells were collected from the plates, washed with distilled water, and resuspended in agroinfiltration buffer (10 mM MES, 10 mM MgCl_2_, pH 5.7). The OD_600_ of the bacterial suspension was measured using a spectrophotometer and adjusted to the desired OD_600_ according to the requirements of the experiment. The prepared bacterial mixtures were incubated with 150 μM acetosyringone for at least one hour in the dark prior to agroinfiltration. For all plant species, the mixtures were infiltrated into the first (most apical) fully-expanded leaf, unless indicated otherwise, using a 1 mL Plastipak syringe without a needle.

### RUBY (betalain) quantification

Betalain extraction and quantification were performed as previously described (Chiang et al., 2024). Briefly, leaf discs from the infiltration spots were collected and incubated in an extraction buffer (10% ethanol, 0.1% formic acid). Betalain was calculated by using a spectrophotometer to obtain the betacyanins (OD_535_) and betaxanthins (OD_475_) values, then subtracting the value measured at OD_600_. For sweet potato, water spinach and common bean, leaf discs were ground in 200 μL of extraction buffer and left under gentle agitation for 10 minutes. Samples were then centrifuged at 4000 rpm for 10 minutes, and 100 μL samples were taken to calculate betalain absorbance.

In faba bean (cultivar ‘de Monica’), betalain accumulation following RUBY expression was quantified using image analysis, as previously described (Chen et al., 2023). Photographs of agroinfiltrated leaves were taken with a Canon EOS 700D camera under standardised settings. Using ImageJ, images were split into RGB channels and the mean pixel density of the agroinfiltrated area was measured in the green channel. The inverse of the green channel measurement was taken as a proxy for pigment intensity.

### Cell death assays

*R. rhizogenes* AS109 containing the required binary plasmid was agroinfiltrated into the abaxial side of faba bean leaves. At 4 days post-infiltration (dpi), leaves were imaged and cell death was recorded under the IR800 CW (800 nm) channel using the Bio-Rad ChemiDoc MP Imaging System. Signal in the 800 nm channel was quantified using the ‘Volume Analysis’ tool in Bio-Rad Image Lab software. The 800 nm signal was normalised to the background or EV signal from the same leaf, depending on the experimental design, and plotted using ggplot2 in R (Wickham et al., 2019).

### Confocal laser scanning microscopy

Confocal microscopy (without infection) was performed on live leaf samples 2-3 dpi. Leaf samples of infiltrated areas were taken using a size 4 cork borer, submerged in distilled water, and mounted on glass slides with glass coverslips. The abaxial side of the leaf discs was imaged using the Leica STELLARIS 1 inverted confocal microscope with a 20x dry or 63x water immersion objective lens. The emission wavelengths for GFP/YFP, BFP and RFP markers were set to 495-550 nm, 402-457 nm and 570-620 nm, respectively. The single-plane confocal micrographs were processed using Leica LAS X software and analysed using ImageJ. To calculate corrected total cell GFP fluorescence, epidermal cells were manually selected to measure integrated density, and neighbouring regions for background fluorescence correction. Corrected total cell GFP fluorescence was calculated for each cell as integrated density minus the area of the cell multiplied by mean fluorescence of the background.

### RNA extraction, cDNA synthesis, and RT-PCR

For RNA extraction, two leaf discs from each of four agroinfiltrated faba bean leaves were taken using a size 4 cork borer. The leaf discs were flash-frozen in liquid nitrogen and total RNA was extracted using the TRIzol RNA Isolation Reagent (Invitrogen) according to the manufacturer’s guidelines. Extracted RNA concentration was measured using a NanoDrop Lite Spectrophotometer (Thermo Fisher Scientific) and 2 μg of RNA underwent DNase treatment using RQ1 ribonuclease-free deoxyribonuclease (Promega). The treated RNA was used as a template for cDNA synthesis by SuperScript IV Reverse Transcriptase (Invitrogen) and subsequent PCR reactions were performed using Q5 High-Fidelity DNA Polymerase (New England Biolabs). Expression of the glyceraldehyde-3-phosphate dehydrogenase (GAPDH) housekeeping gene was used as a transcriptional control (faba bean gene Vfaba.Hedin2.R1.4g150200.1; lettuce gene LOC111886224) (Jayakodi et al., 2023; Reyes-Chin-Wo et al., 2017). The RT-PCRs for GFP expression were performed using primers GFP RT-PCR F7 and GFP RT-PCR R2, for faba bean GAPDH using primers VfGAPDH RT-PCR F3 and VfGAPDH RT-PCR R4, and for lettuce GAPDH using primers Ls_GAPDH_qPCR_F3 and Ls_GAPDH_qPCR_R3 (Supplementary Table 3).

### *Phytophthora palmivora* growth conditions and infection assays

The transgenic *Phytophthora palmivora* LILI KDEL-YFP strain (accession P16830) has been previously described (Rey et al., 2014). *P. palmivora* was cultured on V8 agar plates, supplemented with 100 mg/L G418, at 28°C for 5-7 days before spore collection. Spores were collected by incubating 15 mL of glass-distilled water on a cultured plate for 45 minutes at 22°C. Faba bean leaves were agroinfiltrated 2 days prior to infection. Infections were performed with 10 μL spore droplets on the abaxial side of faba bean leaves and incubated at 22°C for 24-48 hours before imaging by confocal microscopy.

### Data analysis and statistics

All data visualisation and statistics were performed in R version 4.5.2, using ggplot2 and tidyverse packages (Wickham et al., 2019).

## Supplementary Tables

**Supplementary Table 1.** Plant material used in this study.

| Species | Family | Cultivar(s) | Source(s) |
| --- | --- | --- | --- |
| *Petunia hybrida* | Solanaceae | W115 | Academia Sinica transformation core |
| *Nicotiana tabacum* | Solanaceae | Samsun; Xanthi | Kindly donated by Ya-Chun Chang (National Taiwan University) |
| *Solanum tuberosum* | Solanaceae | Kennebec | Local supermarket |
| *S. nigrum* | Solanaceae | NA | Wild accession |
| *Lycium barbarum* | Solanaceae | NA | Local commercial supplier |
| *Physalis peruviana* | Solanaceae | NA | Local supermarket |
| *Ipomoea alba* | Convolvulaceae | NA | Local commercial supplier |
| *I. hederacea* | Convolvulaceae | NA | Wild accession |
| *I. aquatica* | Convolvulaceae | NA | Local commercial supplier |
| *I. batatas* | Convolvulaceae | Taoyuan No. 2 | Local commercial supplier |
| *Mimulus lewisii* | Phrymaceae | LF10 | Kindly donated by Chun-Neng Wang (National Taiwan University) |
| *Salvia splendens* | Lamiaceae | NA | Commercial supplier (Known-You Seed) |
| *Antirrhinum majus* | Plantaginaceae | NA | Commercial supplier (Known-You Seed) |
| *Sesamum indicum* | Pedaliaceae | NA | Local commercial supplier |
| *Coffea canephora* | Rubiaceae | NA | Local commercial supplier |
| *Lactuca sativa* | Asteraceae | NA; Amica | Commercial supplier (Known-You Seed); Commercial supplier (Enza Zaden) |
| *Chenopodium formosanum* | Amaranthaceae | NA | Kindly donated by Na-Sheng Lin (IPMB Academia Sinica) |
| *Brassica rapa* | Brassicaceae | Standard 1-33 | Commercial supplier (Rapid-Cycling Brassica Collection) |
| *Medicago sativa* | Fabaceae | NA | Local commercial supplier |
| *Trigonella foenum-graecum* | Fabaceae | NA | Local commercial supplier |
| *Glycine max* | Fabaceae | Kaohsiung-11 | Local commercial supplier |
| *Pisum sativum* | Fabaceae | Jin Jin | Local commercial supplier |
| *Vicia faba* | Fabaceae | NA; de Monica | Local commercial supplier; commercial supplier (Thompson-Morgan) |
| *Phaseolus vulgaris* | Fabaceae | NA | Commercial supplier (Known-You Seed) |
| *Vigna unguiculata* | Fabaceae | NA | Local commercial supplier |
| *V. radiata* | Fabaceae | Crystal | Kindly donated by Heng-Ming Ting (National Taiwan University) |

**Supplementary Table 2.** Apoplastic elicitors screened for cell death in faba bean.

| Elicitor name | Pathogen | Genbank Gene Accession | Reference |
| --- | --- | --- | --- |
| *Bc*CDI1 | *Botrytis cinerea* | XP_001560184.1 | Franco-Orozco et al. (2017) |
| *Bc*pg1 | *B. cinerea* | XP_001550077.1 | Ji et al. (2023) |
| *Bc*xyn11A | *B. cinerea* | XP_024547454.1 | Frías et al. (2019) |
| *Cc*EG1 | *Cytospora chrysosperma* | ROV90154.1 | Xu et al. (2023) |
| *Cf*ECP2 | *Cladosporium fulvum* | CAA78401.1 | Van den Ackerveken et al. (1993) |
| *Cs*02526 | *Ciboria shiraiana* | WLS32030.1 | Zhang et al. (2024) |
| *Ds*70057 | *Dothistroma septosporum* | EME45906.1 | Hunziker et al. (2021) |
| *Ds*71487 | *D. septosporum* | EME45809.1 | Hunziker et al. (2021) |
| *Ds*74283 | *D. septosporum* | EME40684.1 | Hunziker et al. (2021) |
| *Fg*62 | *Fusarium graminearum* | XP_011320562.1 | Wang et al. (2023) |
| *Gr*EXPB2 | *Globodera rostochiensis* | AJ311902.1 | Ali et al. (2015) |
| *Gr*PEL2 | *G.rostochiensis* | AY094613.1 | Kudla et al. (2007) |
| *Aa*HRIP1 | *Alternaria alternata* | HQ713431.1 | Jones and Perez (2023) |
| *Zt*NLP | *Zymoseptoria tritici* | SMQ56213.1 | Motteram et al. (2009) |
| *Xf*PD_0956 | *Xylella fastidiosa* | AAO28818.1 | Sertedakis et al. (2022) |
| PITG_23123 (SCR50) | *Phytophthora infestans* | XP_002997761.1 | Bos et al. (2003) |
| *Rc*CDI1 | *Rhynchosporium commune* | ARB51385.1 | Franco-Orozco et al. (2017) |
| *Ss*NE1 (SS1G_07027) | *Sclerotinia sclerotiorum* | XP_001591581 | Seifbarghi et al. (2020) |
| SsNE2<br>(SS1G_00849) | *S. sclerotiorum* | XP_001598760 | Seifbarghi et al.<br>(2020) |
| SsXyl2<br>(SS1G_03618) | *S. sclerotiorum* | XP_001595529.1 | Wang et al. (2024) |
| VdEG1<br>(VDAG_04017) | *Verticillium*<br>*dahliae* | XM_009654101.1 | Gui et al. (2017) |
| Zt10<br>(MYCGRDRAFT_11<br>1505) | *Z. tritici* | EGP82940.1 | Kettles et al. (2017) |
| PsXEG1 | *P. sojae* | G4ZHR2.1 | Ma et al. (2015) |

**Supplementary Table 3.**
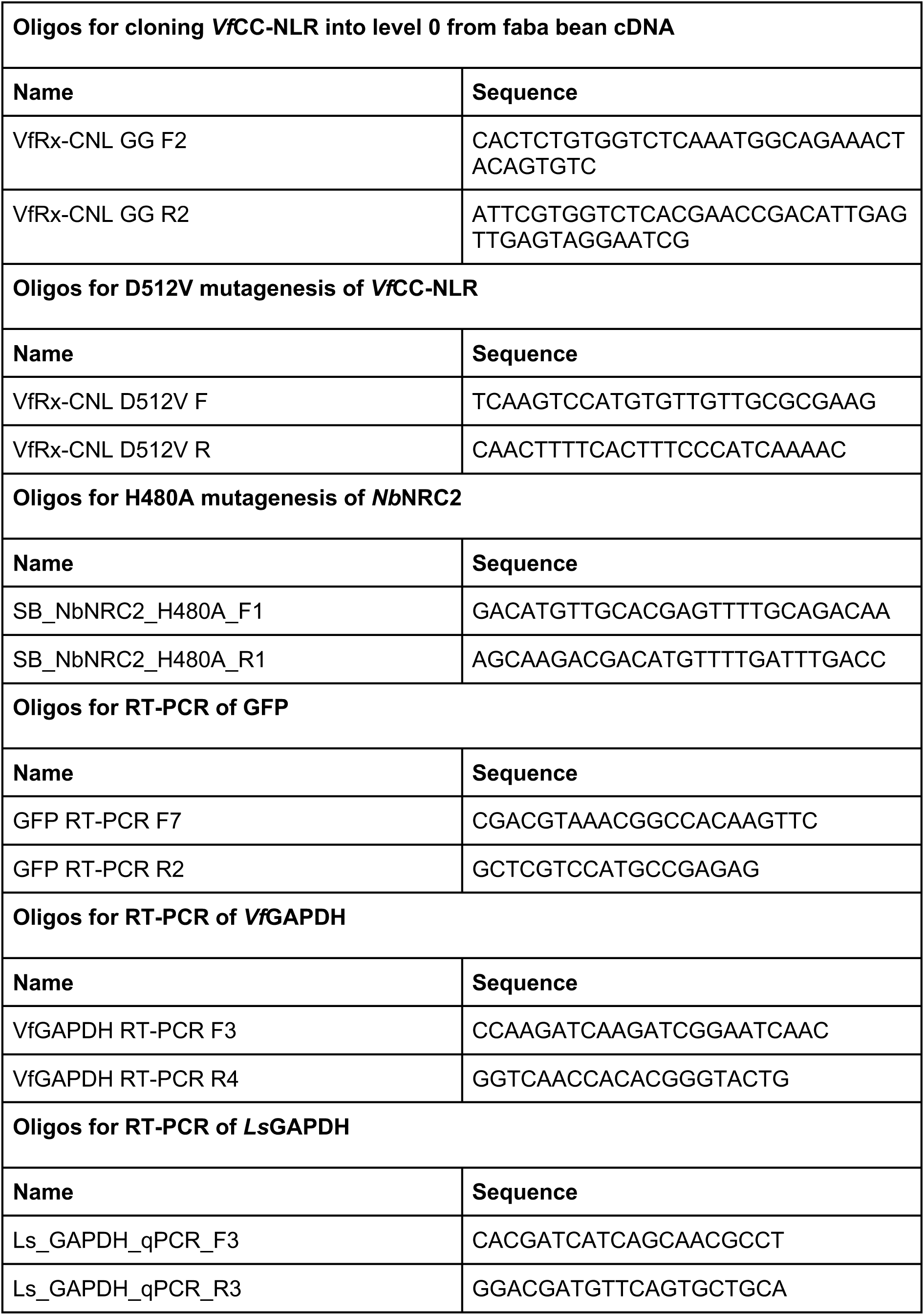
Oligos used for cloning and RT-PCR.

**Supplementary Table 4.**
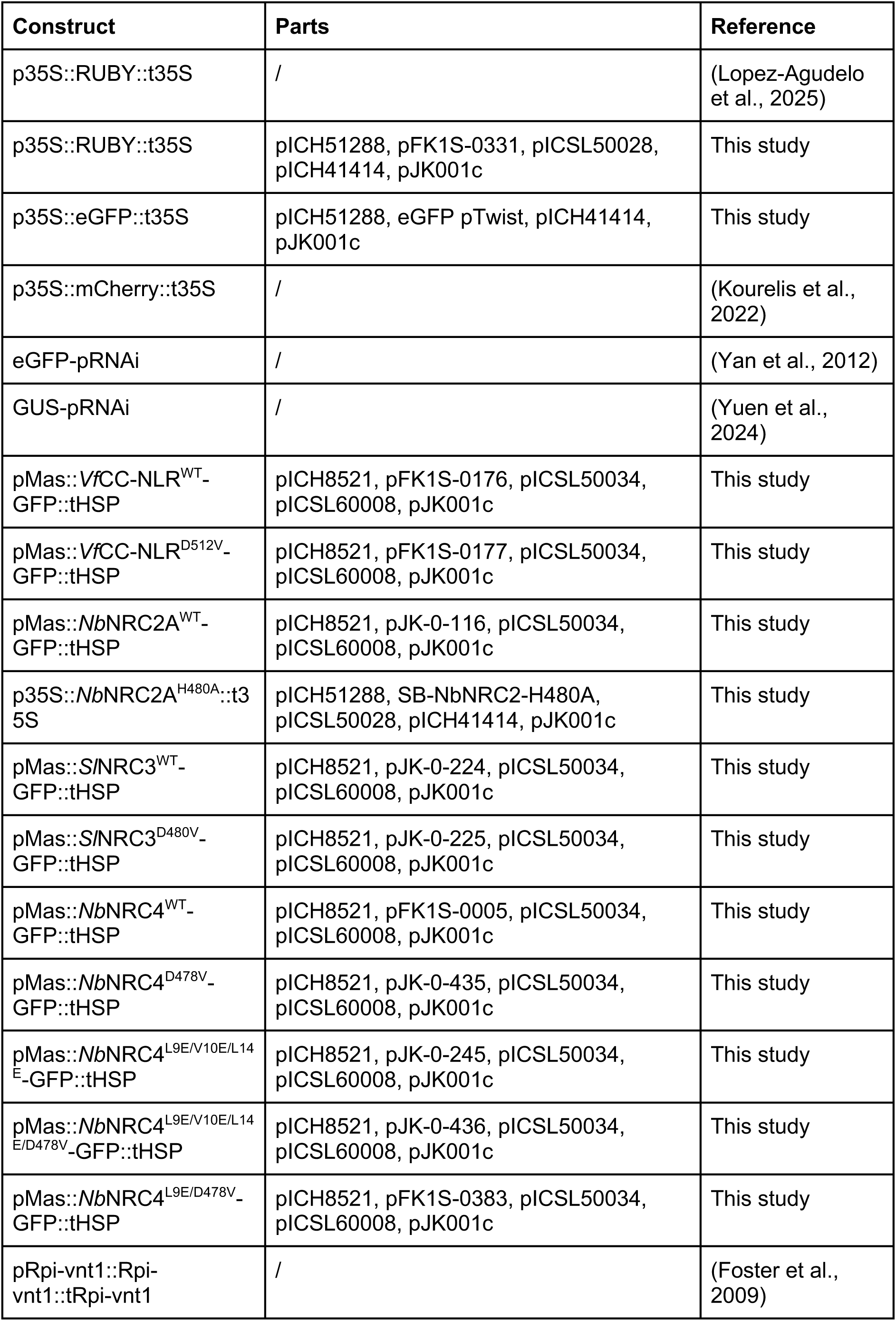
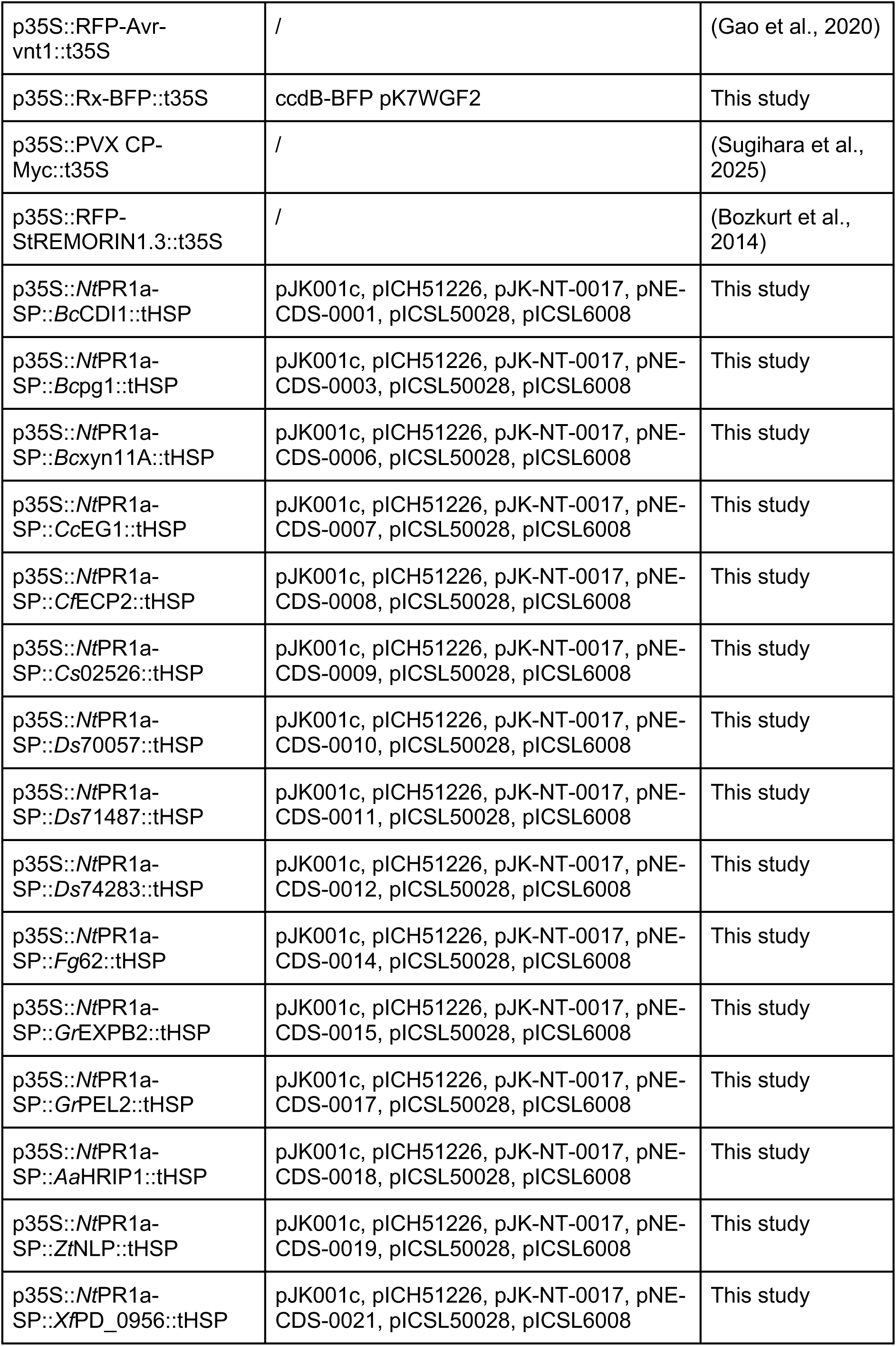
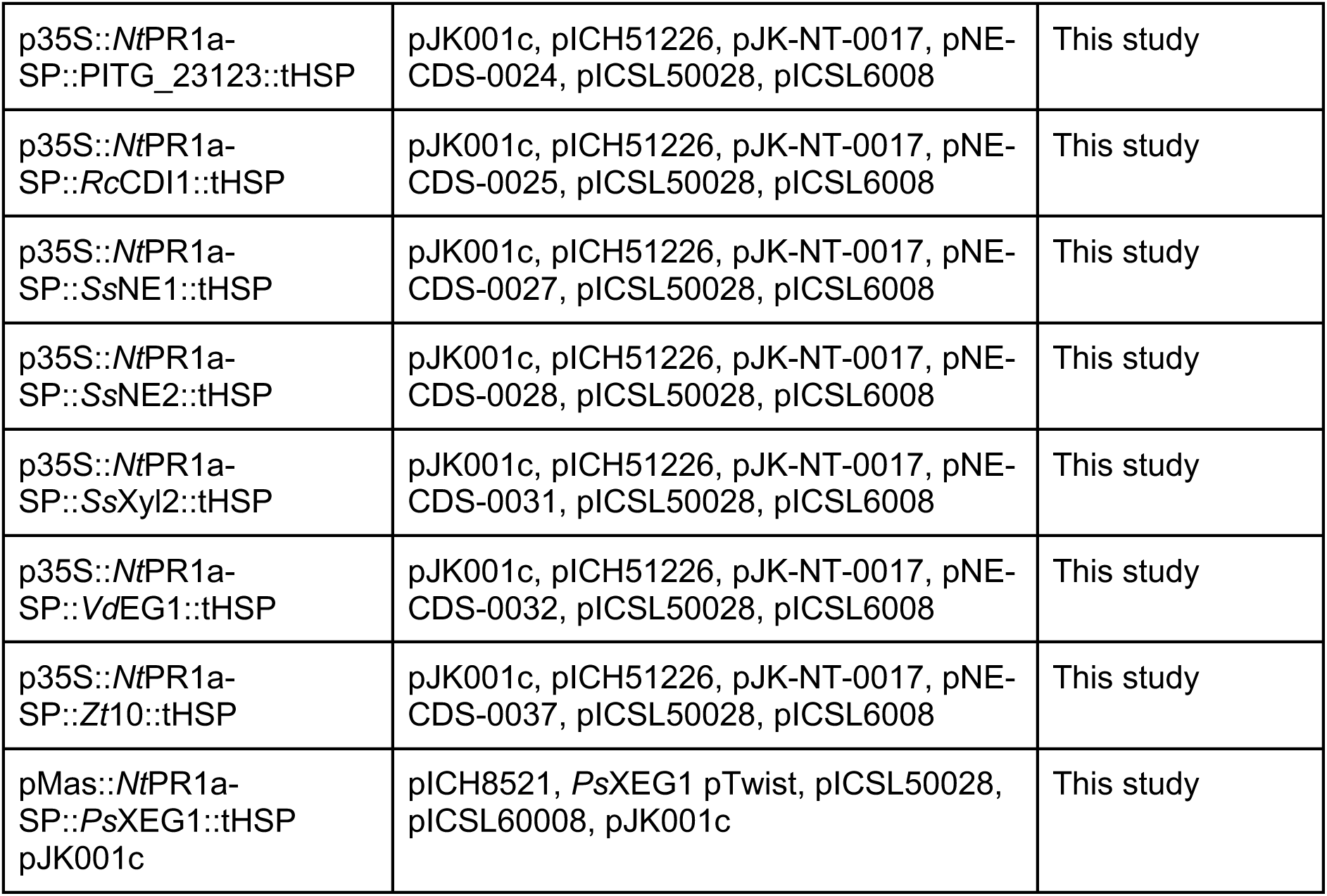
All constructs used in this study.

**Supplementary Table 5.** Details of all *Agrobacterium* and *Rhizobium* strains used in this study.

| Bacterial strain | Genotype | Selection used |
| --- | --- | --- |
| *Agrobacterium fabrum* GV3101 | C58 Rif<sup>R</sup> pMP90 (pTiC58ΔT Gen<sup>R</sup>) | 50 µg/mL rifampicin<br>50 µg/mL gentamycin |
| *A. fabrum* C58C1 | C58 Rif<sup>R</sup> pGV2260 (pTiB6S3ΔT Carb<sup>R</sup>) pCH32<sup>H</sup> | 50 µg/mL rifampicin<br>50 µg/mL carbenicillin |
| *A. fabrum* EHA105 | C58 Rif<sup>R</sup> pEHA105 (pTiBO542ΔT) | 50 µg/mL rifampicin |
| *A. fabacearum* LBA4404 | Ach5 Rif<sup>R</sup> pAL4404ΔT | 50 µg/mL rifampicin |
| *A. fabrum* AGL-1 | C58 Rif<sup>R</sup> Carb<sup>R</sup> pTiBO542ΔT | 50 µg/mL rifampicin<br>50 µg/mL carbenicillin |
| *Rhizobium rhizogenes* AS109 | A4 Rif<sup>R</sup> ΔKan<sup>R</sup> pRiA4ΔT | 50 µg/mL rifampicin |

